# Pseudotyped virus-based platform and structural analysis reveal potential cross-reactivity sites between influenza C and D viruses

**DOI:** 10.64898/2026.07.14.738474

**Authors:** Maria Giovanna Marotta, Shahid Rowles-Khalid, Martin Mayora Neto, Janet M. Daly, Pauline M. van Diemen, Helen E. Everett, Emanuele Montomoli, Claudia Maria Trombetta, Nigel J. Temperton, Kelly da Costa

## Abstract

Influenza C (ICV) and influenza D (IDV) viruses belong to the *Orthomyxoviridae* family and are classified in the genera *Gammainfluenzavirus* and *Deltainfluenzavirus*, respectively. Although the main reservoir of ICV is humans, IDV is mainly found in cattle. To date, the zoonotic potential of IDV has not been fully elucidated. ICV and IDV share about 50% homology at the genetic level, and both express hemagglutinin esterase fusion (HEF) glycoproteins on the surface for the dual purpose of binding the receptor and releasing new virions. Using pseudotyped viruses (PVs) in a pseudotyped virus-based microneutralisation assay (pMN), some bovine serum samples showed strong neutralisation of both ICV and IDV. *In silico* analyses were performed to explore the molecular basis of this phenomenon. HEF structures were recovered from the Protein Data Bank, epitopes were predicted using BepiPred, and sialic acid receptor docking was evaluated with HDOCK. Five potential epitopes were selected, and mutual substitutions of amino acid residues were introduced to generate mutant ICV and IDV HEFs and corresponding PVs. Although only mutant IDV PVs were successfully produced, a reference ICV antiserum showed high neutralising activity against one construct, indicating the exposure of an ICV-like antigenic site within the IDV framework. Herein we provide evidence consistent with the existence of antigenic sites shared between ICV and IDV, which could be exploited for cross-protective vaccine design, through integrated computational and experimental investigations.

## Introduction

Influenza C (ICV) and influenza D (IDV) viruses are members of the *Orthomyxoviridae* family, classified as *Gammainfluenzavirus* and *Deltainfluenzavirus* genera, respectively. Unlike influenza A (IAV) and B (IBV) viruses with genomes consisting of eight segments and encoding two glycoproteins, hemagglutinin (HA) and neuraminidase (NA), ICV and IDV incorporate seven genome segments and only one major envelope glycoprotein, hemagglutinin esterase fusion (HEF) (1). HEF combines the functions of both HA and NA, being responsible for receptor binding, receptor destruction through its esterase activity, and membrane fusion (2). Whilst the impact on public health by IAV and IBV is well characterised, ICV and IDV have received less attention despite their increasing relevance. ICV was first isolated in 1947 from a human sample during an epidemic of respiratory illness (3). Primarily infecting children under two years of age (4), ICV has also been reported to infect animals, such as pigs and cattle, with serological evidence also found in dogs (5, 6, 7, 8). IDV was isolated from pigs in the USA in 2011 and later from cattle, which were found to be the primary livestock reservoir (9, 10). Subsequent serological and molecular surveillance studies have demonstrated widespread circulation of IDV in bovine populations across multiple geographic regions, including Europe and Asia, reflecting growing global interest in this virus and its epidemiology (11, 12, 13). Importantly, IDV has been recognised as a contributing pathogen in bovine respiratory disease (BRD), a multifactorial syndrome that represents one of the most prevalent and costly diseases in cattle (14). Although IDV has not been associated with human illness, seropositivity has been observed, particularly in individuals with occupational exposure (15, 16), and growing evidence from experimental models and human surveillance studies suggest a potential zoonotic risk (17, 18).

Despite ICV and IDV belonging to different genera, their HEF glycoproteins share approximately 53% amino acid homology, and their 3D structures are highly conserved (19). Both viruses recognise N-acetyl-9-O-acetylneuraminic acid (Neu5,9Ac) through the HEF glycoprotein, but while ICV has a restrictive receptor binding domain (RBD) and a conserved glycosylation site near the RBD, that if occupied might restrict the accessibility to the receptor (2, 19, 20, 21), IDV has a salt bridge and glycosylation sites that can influence the tropism of the virus and result in a more open receptor binding channel (19). This structural configuration allows IDV to interact more efficiently with a wider range of host cells, compared to ICV.

As the only envelope glycoprotein, HEF represents an important target for neutralising antibodies (19). The preservation of structural features of HEF between species leads to the hypothesis that cross-reactive antibodies may be produced during infection with either ICV or IDV. To our knowledge, this has not been addressed experimentally despite both viruses having been isolated from cattle showing respiratory signs consistent with BRD, hinting at a co-pathogenic relationship (22).

Computational analysis is a useful tool to exploit existing antigenic structures for the design of novel vaccine strategies or targets. These approaches have been employed as complementary tools in influenza virus and vaccine research (23, 24). To explain serological observations, we have applied a computational pipeline to further characterise the HEF glycoproteins of ICV and IDV, analysing their 3D structures, simulating docking interactions with receptors and predicting antigenic epitopes. Predicted epitopes were subsequently mapped onto the HEF structural models to identify conserved and distinct antigenic regions. This insight informed the design of plasmid constructs carrying targeted point mutations, which were used to produce PVs incorporating the mutated HEF glycoproteins, enabling the functional assessment of antibody recognition. Using this novel approach of combined *in silico* and *in vitro* techniques, we identified potential antigenic epitopes shared between ICV and IDV HEFs and evaluated their contribution to antibody binding, providing new insights into the conserved molecular determinants among these viruses. Although the role of these epitopes in mediating cross-protective immunity has not been established yet, our findings have important implications for understanding antigenic evolution, improving zoonotic surveillance, and guiding the rational design of next-generation influenza vaccines.

## Methods

### Samples and antisera

A total of 46 cow serum samples were collected across the UK between 2012 and 2013. Sera were heat-inactivated at 56°C for 30 minutes prior to use. To assess cross-reactivity with other influenza subtypes, rooster antiserum to C/Taylor/1233/1947 (NR-3132, BEI Resources, NIAID, NIH) reconstituted with 500 *µ*L of sterile dH_2_O was used as a positive control for ICV, while a serum sample from a pig vaccinated against D/Swine/Oklahoma/1334/2011 was used as a positive control for IDV (25). Commercial antisera obtained from NIBSC were tested against IC_Minnesota_- and ID_Italy_-PVs for reciprocal cross-reactivity assessment. These included sheep antisera retrieved to A/Puerto Rico/8/34 (H1N1) (NIBSC, code 03/242), A/Darwin/9/2021 (H3N2) (NIBSC, code 22/226), B/Brisbane/60/2008 (Victoria lineage) (NIBSC, code 16/192), and B/Phuket/3073/2013 (Yamagata lineage) (NIBSC, code 17/214).

### Cell lines

Human embryonic kidney 293T/17 (HEK293 T/17, ATCC: CRL-11268a) cells were used for transfection. Swine testicular (ST, ATCC: CRL-1746) cells were used as target for transduction efficiency of mutants PVs. Both the cell lines were cultured in complete Dulbecco’s Modified Essential Medium (DMEM) (PANBiotech, P04-04510), with high glucose and GlutaMAX supplemented with 10% (v/v) heat-inactivated fetal bovine serum (PANBiotech, P30-8500) and 1% (v/v) penicillin–streptomycin (PenStrep) (Sigma, P4333) at 37°C with 5% CO_2_.

### Structural analysis of HEF glycoproteins

To investigate the molecular basis underlying the potential cross-reactivity between ICV and IDV, the available crystallographic structures of HEF glycoproteins were retrieved from the Protein Data Bank (PDB) (26): C/Johannesburg/1/66 (PDB ID: 1FLC) and of D/Swine/Oklahoma/1334/2011 (PDB ID: 5E64). Since these HEF structures differ from the PV strains used in functional assays (C/Minnesota/33/2015 and D/Swine/Italy/199724-3/2015), pairwise sequence alignments were performed using NCBI Protein BLAST to assess identity and conservation (27). To predict potential epitopes, full-length amino acid sequences of HEFs in FASTA format were submitted on BepiPred version 3.0 epitope prediction tool (28).In this context, these amino acids are incorporated into the protein chain order and are referred to as residues, reflecting their covalent linkage via peptide bonds. Predicted epitopes were mapped on 3D models of HEFs and visualised using PyMOL version 3.0 (Schrödinger, LLC) (29). In addition, to explore HEF interaction with sialic acids, molecular docking was performed using the HDOCK web server (30).

### Design of mutants for HEF glycoproteins and plasmid transformation

Residues located within predicted epitopes of ICV and IDV were selected for mutual substitution to investigate their potential role in antibody recognition. Three HEF mutant constructs were generated for ICV (HEF_Minnesota_-MUT1, -MUT2 and -MUT3) and two for IDV (HEF_Italy_-MUT1 and -MUT2). All constructs were synthesised and subcloned into the mammalian expression plasmid pcDNA3.1+ by GenScript (UK). Plasmid transformation was performed via the heat-shock method in chemically induced competent *E. coli* XL10-gold Ultracompetent cells (Agilent 200315) (ICV), and *E. coli* DH5*α* cells (Invitrogen 18265-017) (IDV). Plasmid DNA was recovered from transformed bacterial cultures incubated at 33°C (ICV) and at 37°C (IDV), via the plasmid mini kit (Qiagen 12125, Manchester, UK) and quantified using UV spectrophotometry (NanoDrop™—Thermo Scientific, Paisley, UK) (25).

### Pseudotyped virus (PV) production, titration and pseudotyped virus-based microneutralisation (pMN) assay

PVs were produced and titrated following the optimised protocol (31). Titres were measured as relative luminescence units (RLU/mL) after transduction of ST cells. PVs incorporating HEF glycoproteins from ICV (C/Minnesota/33/2015) and IDV (D/Swine/Italy/199724-3/2015) were used in a pMN assay, alongside PVs representing seasonal human influenza strains (32) including A/England/195/2009 (H1), A/Switzerland/7515293/2013 (H3), B/Washington/2/2019 (Victoria lineage), and B/Phuket/3073/2013 (Yamagata lineage), to assess cross-reactivity. The pMN assay was performed as previously described (25). Briefly, cow serum samples were diluted 1:40, and ICV and IDV antisera were diluted 1:100. A serial two-fold dilution was performed for all the samples and antisera before adding ICV/IDV PVs at a concentration of 2 × 10^7^ RLU/mL. After incubating for an hour at 37°C, 1.75 × 10^4^ cells/well of ST cells were added before plates were incubated for a further 48 hours at 37°C with 5% CO_2_. After this period, all media was removed and 25 *µ*L of Bright-Glo™ Luciferase Assay System (Promega, Southampton, UK) diluted 1:1 with PBS was added to each well, and read using a GloMax® Navigator luminometer (Promega, Southampton, UK) with the Promega GloMax® Luminescence Quick-Read protocol. Serum IC_50_ titre is expressed as the reciprocal of the dilution resulting in 50% reduction of the pMN readout, with a higher value indicating improved neutralisation activity.

### Statistical analysis

All statistical analysis was performed with GraphPad Prism Version 10.3.1 (GraphPad Software). PV titres were estimated using Microsoft Office Excel 365 (Version 16.92, Microsoft Corporation, Redmond, WA, USA). Titres from pMN assays were normalised to cell only (100% neutralisation) and PV only (0% neutralisation) controls. IC_50_ values were calculated by a non-linear regression model (log [inhibitor]) vs. normalised response-variable slope) (31). An unpaired two-tailed Student’s test analysis was performed to calculate statistical significance (*, **, ***, and **** = *p* < 0.05, significant differences; ns = not significant).

## Results

The neutralising activity of 46 cow serum samples was measured against IC_Minnesota_- and ID_Italy_- PVs. Neutralising titres against IC_Minnesota_ ranged from 28.6 to 8712 (median IC_50_ = 1337.5), whereas titres against ID_Italy_ ranged from 40 to 5062 (median IC_50_ = 347). In addition, 54.3% (25/46) of sera showed IC_50_ titres ≥1000 against IC_Minnesota_ compared with 28.3% (13/46) against ID_Italy_. This difference was statistically significant, indicating a markedly stronger neutralising response to the ICV PV (Figure 1A). This prompted us to explore whether the positivity observed was the result of cross-reactivity between the two viruses. To address this, the ability of subtype specific antisera (anti-IAV/IBV/ICV/IDV) to neutralise IC_Minnesota_- (Figure 1B) and ID_Italy_-PVs (Figure 1C) was assessed. All heterologous antisera failed to achieve 50% neutralisation against IC_Minnesota_-PV (Figure 1B). Neutralisation by mismatched antisera generally remained below 30%; however, the IDV antiserum reached approximately 45% of neutralisation at its highest activity. In contrast, the ICV antiserum cross achieved >50% neutralisation against ID_Italy_-PV (Figure 1C), showing an asymmetrical cross-neutralisation pattern and highlighting the need for further investigation. The remaining heterologous antisera showed neutralisation levels below approximately 30%. To eliminate background or non-specific neutralisation across influenza viruses, both ICV and IDV antisera were tested against IAV H1N1 and H3N2, and IBV Victoria and Yamagata lineages PVs (Figures 1D, E, F & G, respectively). Minimal neutralising activity that remained below the 50% of neutralisation threshold was observed, with peak neutralisation values of approximately 45% for H1N1 and 25% for H3N2 (Figures 1D & E). A higher cross-reactivity was observed against both IBV PVs, particularly against the Yamagata lineage, where the ICV antiserum reached approximately 70% neutralisation and the IDV antiserum reached approximately 85% neutralisation (Figures 1F & G).

**Figure 1.**
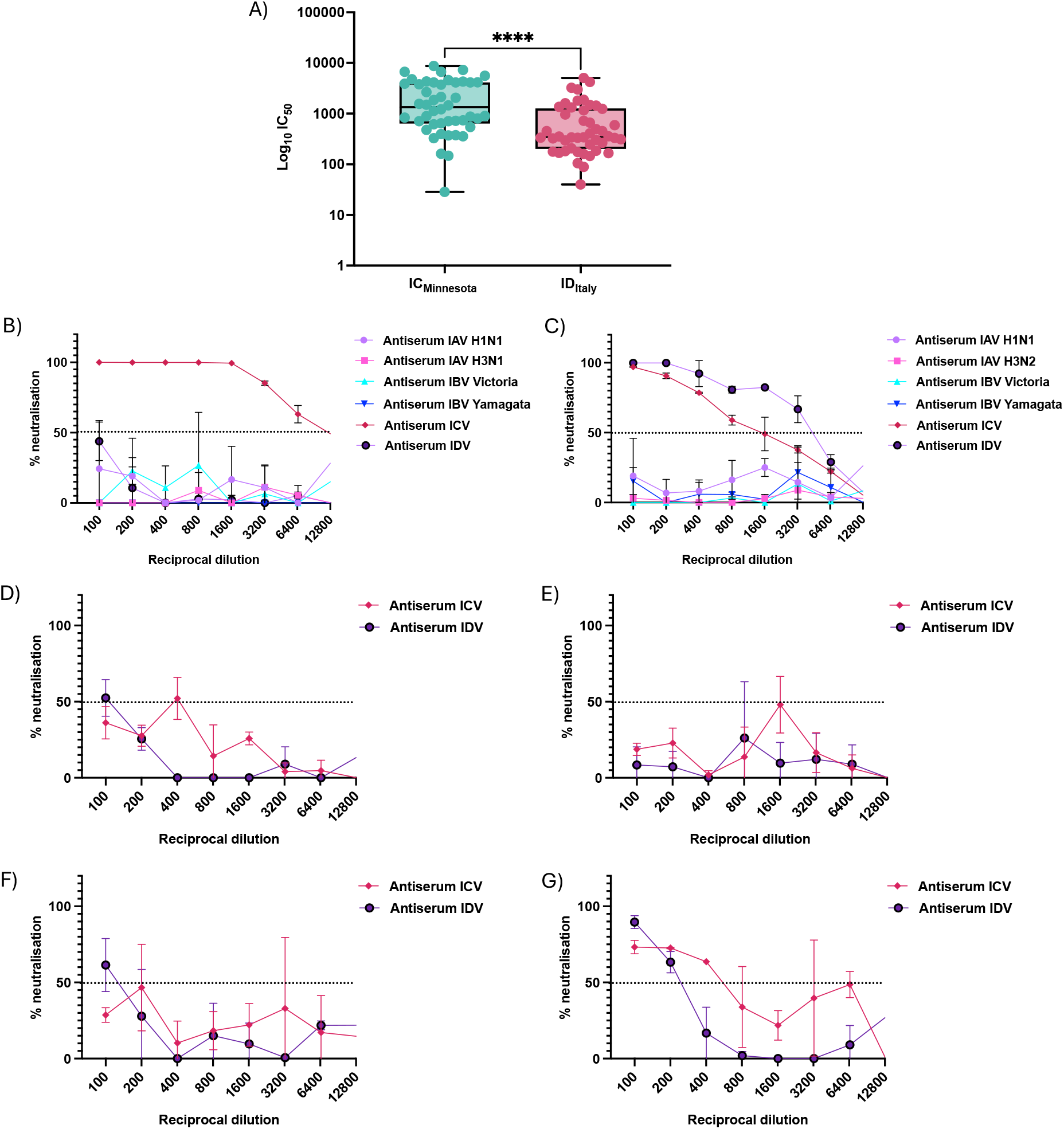
A) Neutralisation titres of cow serum samples versus IC_Minnesota_- and ID_Italy_-PVs. Titres are expressed as IC_50_, plotted on a log_10_ scale. Boxes indicate the central distribution of the data, the horizontal lines represent the median, and the whiskers show the spread of the values. Statistical significance: *p* < 0.05 (****). Neutralisation curves of B) IC_Minnesota_- and C) ID_Italy_-PVs by homologous ICV and IDV antisera as well as by heterologous antisera to IAV (H1N1, H3N2) and IBV (Victoria and Yamagata lineages). Neutralisation curves of D) IAV (H1N1), E) IAV (H3N2), F) IBV (Victoria lineage) and G) IBV (Yamagata lineage) PVs by ICV and IDV antisera. Error bars represent ± standard deviation of two replicates per dilution.

To better define the molecular basis of potential cross-reactivity between ICV and IDV, we aligned the HEF glycoproteins expressed on PVs used here (Figure 1A) with the corresponding reference HEF sequences available in the databases for computational investigation. Amino acid sequence alignment confirmed a high degree of conservation with 96.79% identity for ICV and 98.19% for IDV (Figure S1). To predict the potential cross-reactive epitopes of HEF, amino acid sequences were analysed using BepiPred. Predicted epitopes with scores exceeding the default threshold of 0.15 were selected for further investigation. The threshold of 0.15 was chosen based on the performance of the prediction model during its development and validation to represent a balanced compromise between sensitivity (true positive rate) and specificity (true negative rate), maximising correct predictions of epitope residues while minimising false positives. A total of five epitopes were predicted for both ICV and IDV (Figures 2A & B, respectively).

**Figure 2.**
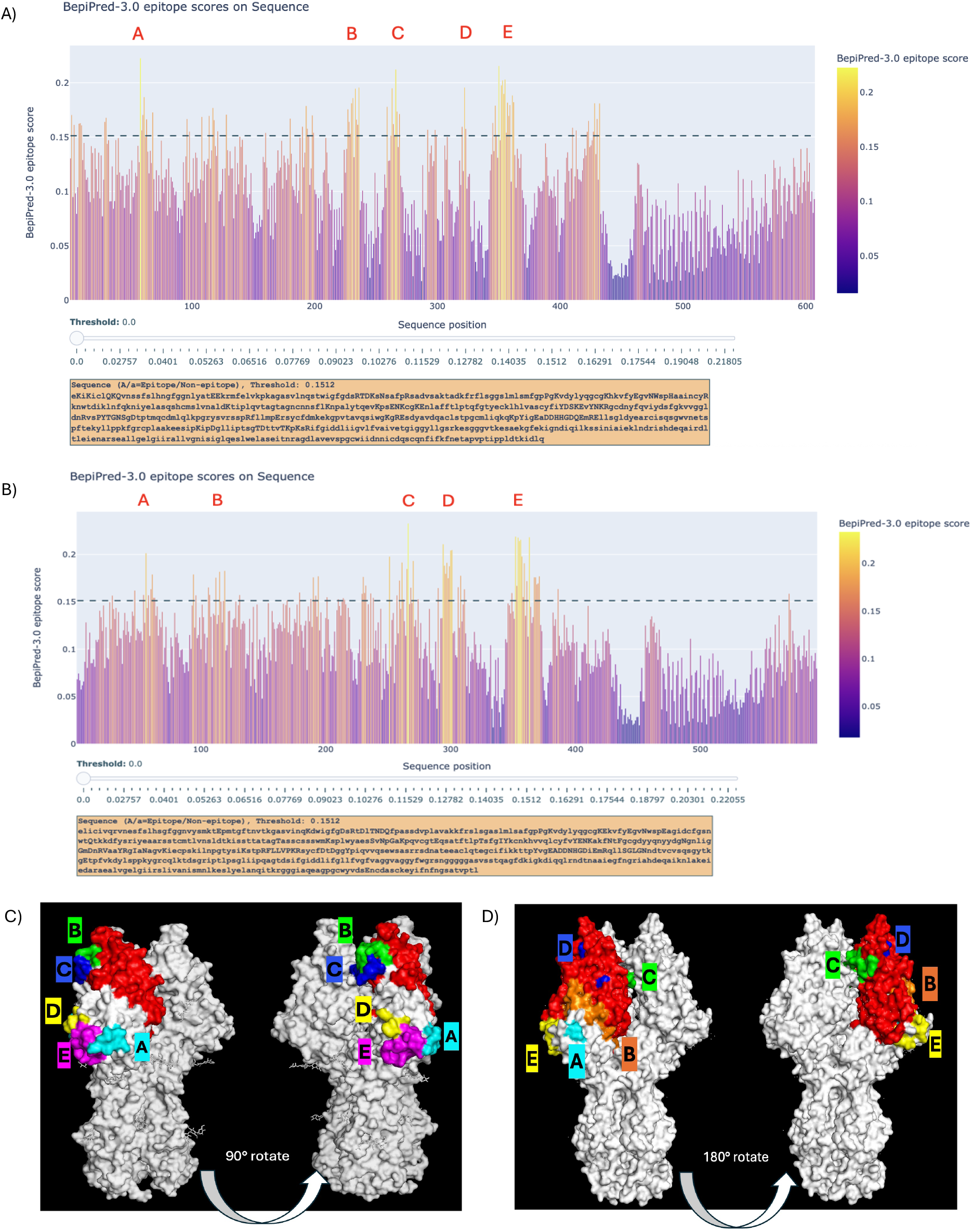
Map of the HEF predicted epitopes of A) ICV and B) IDV using BepiPred. C-D) Structural mapping of predicted epitopes (A–E) to a single monomer within the trimer of C) ICV HEF and D) IDV HEF using PyMOL. Each epitope is highlighted in a distinct colour and displayed in two orientations allowing a clear visualisation of their spatial localisation. The receptor binding domain (RBD) is represented in red.

Structural visualisation of predicted epitopes using PyMOL was performed with surface and cartoon-chain visualisation settings to highlight the 3D topology of HEF proteins. Residue numbering was maintained in accordance with the sequence format used in PDB reference structures and BepiPred input files to ensure consistency. This analysis showed that many of these regions are surface-exposed and mapped to or near the RBD (highlighted in red), of ICV (Figure 2C) and IDV (Figure 2D). Their localisation and accessibility support the hypothesis that these epitopes may act as key determinants of antibody recognition between the two viruses.

To not interfere with recognition of the host receptor, we intentionally selected epitopes outside the RBD of the HEF of both ICV and IDV for mutant constructions. Epitopes A (cyan), D (yellow) and E (magenta) were selected for ICV, and epitopes A (cyan) and E (yellow) were chosen for IDV (Figure 2). To further ensure these regions were not occluded by receptor binding, molecular docking predictions for ICV and IDV HEFs interaction with the two known sialic acid receptors, Neu5Ac and Neu5,9Ac, was performed. In ICV, Neu5Ac was predicted to bind within the HEF2 region, on the inner side of the trimer and less exposed to the host protein surface (Supplementary figure S1A), whereas Neu5,9Ac instead bound to an upper and more exposed region of the protein (Supplementary Figure S1B). A similar situation was obtained for IDV, with rather Neu5Ac bound to the top of the glycoproteins (Supplementary figure S1C) whereas Neu5,9Ac interacted in a proximal but different site of the same exposed surface (Supplementary figure S1D). Significantly, docking analyses suggested that the selected epitope regions did not spatially overlap with the predicted receptor interaction sites, supporting their selection for mutagenesis without definitively excluding potential indirect structural or functional effects. Structural mapping subsequently indicated that these regions were positioned within the esterase domain of HEFs (data not shown).

Based on the alignment of HEF glycoproteins from ICV and IDV, mutants were generated by specific residue substitutions between the two sequences at the highlighted homologous positions (Supplementary figure S2). This strategy resulted in the generation of three ICV mutants (HEF_Minnesota_ MUT1–3) carrying substitutions at epitopes A, D and E, and two IDV mutants (HEF_Italy_ MUT1–2), carrying substitutions at epitopes A and E (Supplementary table S1). In this way, the residues of the two viruses were exchanged at the same structural positions by not changing the general structure of the HEF glycoproteins.

The mutant HEF-encoding plasmids were transformed for PV production. IC_Minnesota_-HEF mutants did not generate high-titre PVs; however, low but measurable PV titres (∼10^6^ RLU/mL) were detected (Supplementary figure S3A). In contrast, ID_Italy_-HEF mutant PVs were successfully generated, achieving high titres (>10^8^ RLU/mL) (Supplementary figure S3B). To investigate the potential structural effects associated with the introduced mutations, a computational structural investigation of epitopes was performed on PyMol. Mutations were mapped onto the 3D structure of HEFs and compared with corresponding wild-type models. For ICV HEF mutants, this analysis revealed localised structural alterations compared to the wild-type glycoprotein (Figure 3A). In contrast, no major structural changes were observed in the corresponding IDV HEF mutant models (Figure 3B).

**Figure 3.**
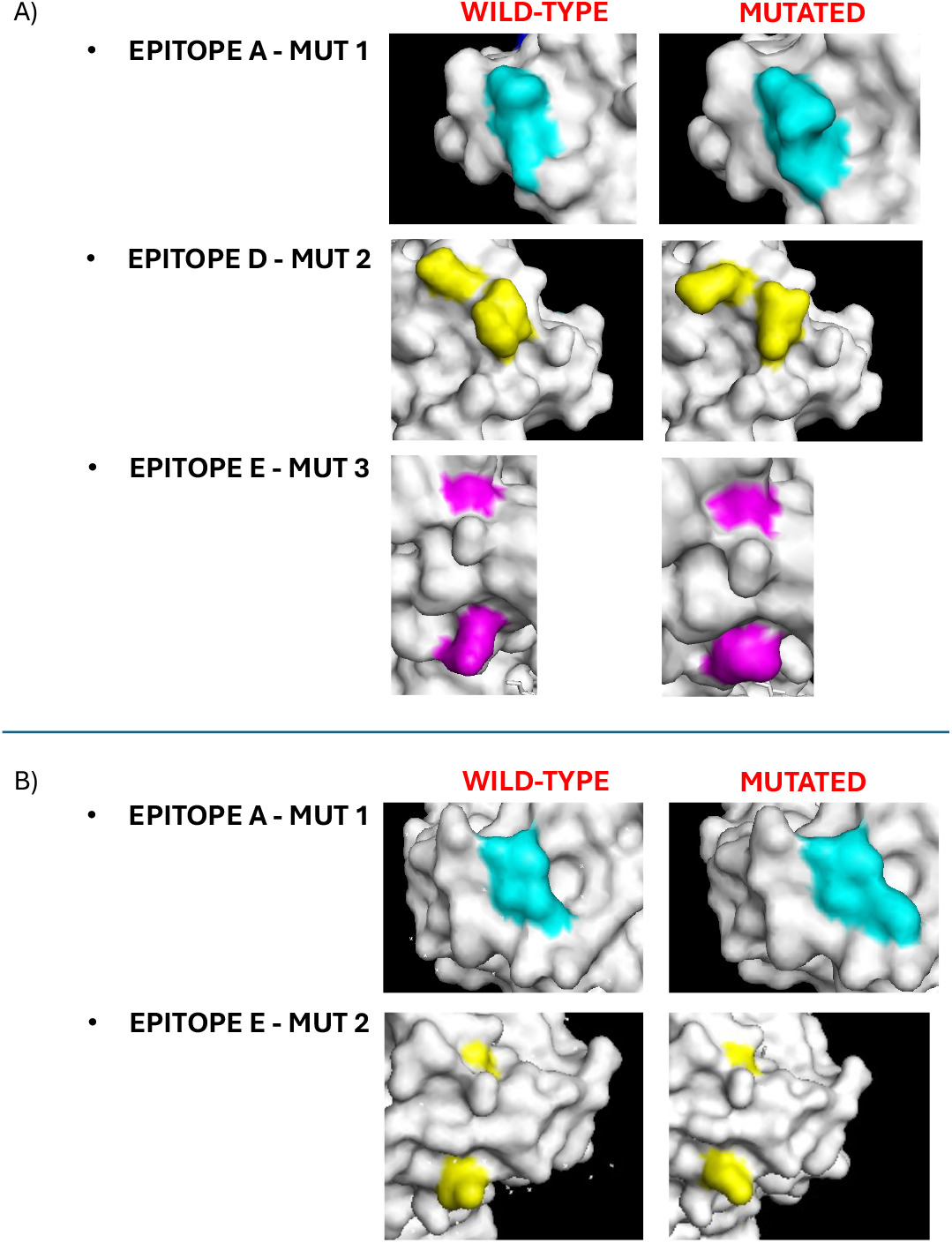
Structural mapping of wild-type and corresponding mutant HEF epitopes of A) ICV and B) IDV is visualised using PyMOL. Structures are displayed in the same orientation to allow direct structural comparison.

ICV and IDV antisera were then assessed for pMN activity using the ID_Italy_-mutant PVs. Using ICV antiserum, ID_Italy_-MUT1 was neutralised more effectively than both ID_Italy_-MUT2 and ID_Italy_ wild-type PVs, providing further evidence that cross-reactivity was enhanced (Table 1). In contrast, ID_Italy_-MUT2 had no appreciable difference in neutralisation compared to ID_Italy_ wild-type PV (Table 1). Conversely, the IDV antiserum showed robust and comparable neutralising activity against all PVs tested without significant differences in IC_50_ titres (Table 1, Supplementary figure S3C&D).

**Table 1.** Comparison of neutralisation activity (IC_50_ titre) of ICV and IDV antisera tested using ID_Italy_- mutant and wild-type PVs.

| PVs | ICV antiserum (IC<sub>50</sub>) | IDV antiserum (IC<sub>50</sub>) |
| --- | --- | --- |
| ID<sub>Italy</sub> wild type | 807 | 3712 |
| ID<sub>Italy</sub>-MUT1 | 1564 | 3646 |
| ID<sub>Italy</sub>-MUT2 | 847 | 3913 |

## Discussion

This study was initiated following the observation that the neutralising activity of cow serum samples was higher against ICV than IDV PVs. Additionally, this phenomenon was not observed when evaluating neutralising activity against IAV or IBV, although some background activity was likely due to the polyclonal nature of the antisera and possible recognition of conserved carbohydrate groups.

Given the sequence and structural similarity between ICV and IDV HEFs, we hypothesised that some of these antigenic regions might be functionally interchangeable, resulting in cross-reactivity and potentially contributing to the increased neutralisation observed when measured using ICV PVs. Interestingly, an asymmetric neutralisation pattern was observed, whereby ICV antiserum appeared to neutralise IDV more efficiently than the reciprocal condition. This asymmetry may indicate that antigenic determinants of the two viruses are differentially targeted during infection. Herein, we have provided insights into the basis of this observation, raising the possibility that prior exposure to ICV could generate antibodies capable of recognising related antigenic regions on IDV, and vice versa.

An epitope exchange approach was adopted, in which selected ICV epitopes were introduced into the IDV HEF sequence, while reciprocal substitutions were also evaluated in the ICV HEF backbone. This combined bioinformatic and experimental approach enabled the assessment of epitope mutations while attempting to minimise disruption to protein folding or function, increasing the chances of experimental success and improving the predictive power of computer modelling. Comparison of predicted regions revealed that some epitopes were conserved between the two viruses, suggesting the presence of shared antigenic characteristics. The docking simulations with sialic acid receptors confirmed distinct binding patterns between ICV and IDV. Importantly, these receptor-binding regions were deliberately excluded from mutagenesis to minimise disruption of receptor recognition and viral entry, allowing the study to focus on the antigenic role of selected non-RBD epitopes. The primary interaction of ICV with Neu5,9Ac was localised to the upper portion of the HEF1 subunit, consistent with its limited receptor specificity (19). In contrast, IDV bound to both Neu5,9Ac and Neu5Ac in the most exposed areas of HEF1, supporting previous observations that IDV can involve multiple sialic acid receptors and infect a wider range of hosts (19, 33). HDOCK uses a hybrid docking strategy that integrates both model-based and *ab initio* approaches. This combination allows unbiased research across the entire protein surface rather than limiting anchoring to predefined binding regions. As a result, HDOCK can identify potential interaction sites even in the absence of prior structural information. However, model-based docking depends on the availability and quality of suitable structural models, whereas *ab initio* docking, while more flexible, tends to be computationally demanding and may produce less accurate predictions. Future studies could therefore build on these exploratory analyses using more biologically guided docking platforms, such as HADDOCK, to refine the predicted interaction interfaces.

Only ID_Italy_-mutant PVs were successfully produced, which could be explained by our computational analyses which revealed localised conformational alterations in ICV HEF mutant models. Although these *in silico* observations do not establish a direct causal relationship, they are consistent with the hypothesis that specific substitutions may differentially influence the structural stability of the glycoprotein. The selected epitopes are located within regions of the esterase domain but remain distinct from the catalytic triad responsible for esterase activity Ser57, Asp356, His359 (19). Evaluation of HEF esterase activity would be necessary to exclude potential indirect effects on enzymatic function. It is also possible that the introduced substitutions influenced the glycoprotein folding or maturation. In this context, evaluating HEF expression levels by Western Blot and incorporation into PVs would help clarify whether the reduced titres reflect these effects.

Functional analysis of the ID_Italy_-mutant PVs using homologous and heterologous antisera revealed that ICV antiserum exhibited increased neutralising titre activity to the ID_Italy_-MUT1 construct. The finding implies that the single residue substitution introduced may have exposed or altered an antigenic determinant recognised by antibodies raised against ICV, leaving the epitopes targeted by IDV antibodies unaffected. Importantly, this result is consistent with the asymmetric cross-neutralisation pattern initially observed, further suggesting that shared antigenic determinants may be more accessible or more effectively recognised within the IDV HEF glycoprotein. These findings provide preliminary evidence that specific shared HEF regions can accommodate substitutions without completely disrupting antigenic recognition and offer insight into the structural flexibility of IDV. However, differential structural tolerance between different HEF isoforms would require the evaluation of additional mutations in multiple regions of the glycoprotein. A broader investigation of shared and distinct epitopes across multiple HEF domains will be needed to determine whether this structural tolerance extends beyond the sites examined in this study. Additionally, precise attribution of neutralisation to individual residues will require detailed epitope mapping and characterisation of monoclonal antibodies.

Recent antigenic characterisation studies of IDV HEF have identified both lineage-specific and conserved neutralising epitopes, supporting the concept that structurally conserved HEF regions could be exploited for the design of broadly reactive immunogens (34). In this context, our epitope exchange strategy provides a preliminary framework to explore the feasibility of engineered HEF immunogens aimed at enhancing cross-reactive antibody responses. Although conserved HEF regions were identified through structural analyses, the potential influence of glycosylation on epitope accessibility and antibody recognition requires further investigation.

The zoonotic potential of IDV has recently become a topic of scientific discussion following the detection of bovine-adapted avian IAV H5N1 and its transmission to humans (35). This is especially important given that the main livestock reservoir of IDV is cattle (9, 10). PCR-based studies have provided evidence of ICV infection in cattle with respiratory disease in the USA, which supports the possibility that both ICV and IDV may co-circulate within the same host population (6). Additionally, there have been reports of human exposure, further complicating the immunological landscape (17, 18). However, whilst the increase in sero-surveillance is encouraging (16), our understanding of IDV biology, transmission and immunity of IDV remains limited compared to the extensive knowledge accumulated for IAV and IBV.

The envelope glycoprotein HEF of ICV and IDV, share approximately 53% amino acid identity (19). This genetic similarity raises the possibility of cross-reactive immune responses between the two viruses. Furthermore, the potential existence of cross-protective immunity may impact the disease outcome in shared hosts, such as cattle with possible implications for zoonotic transmission. In support of this possibility, the asymmetric neutralisation pattern observed in this study demonstrated that ICV-specific antisera neutralised IDV more efficiently than the reciprocal case, suggesting that prior exposure to ICV may partially trigger immune recognition of antigenically related regions within the IDV HEF glycoprotein. Although the extent of any protective effect remains unclear, such immune priming could potentially modulate infection outcome, transmission dynamics, or serological interpretation in both humans and bovine populations where these viruses can co-circulate. In humans, most individuals develop antibodies against ICV during childhood, but these antibody responses tend to wane into adulthood (36). Further investigations are required to assess potential effects on infection susceptibility by antigenically related viruses such as IDV.

This study illustrates the importance of hybrid computational-experimental analysis to characterise less studied viruses, with limited experimental data and reagents available, as well as the use of novel approaches, such as use of the high-throughput PV platform (25, 31, 32). Modelling and evaluating *in silico* viral wild-type and mutant structures prior to laboratory experimentation provides predictive information for optimisation and experimental design prior to testing *in vitro*. Nevertheless, some limitations are recognised, such as the use of polyclonal antisera that complicates the ability to determine the neutralisation effects to specific epitopes, and the absence of monoclonal antibody validation. The computational analyses were restricted to linear epitopes, leaving conformational determinants unaddressed. Another limitation concerns the absence of negative controls in docking experiments such as non-interacting protein pairs, that would have provided an internal benchmark for docking specificity. Integrating such controls into future studies could improve the interpretability of docking-based predictions.

ICV and IDV research findings highlight the interconnectedness of human, and animal health and importance of improved understanding (37). The circulation of IDV in cattle and the detection of neutralising antibodies in humans underscore the relevance of monitoring antigenic overlap across species. Data presented here confirm the structural conservation of HEF and support the existence of antigenic overlap between ICV and IDV. In the future, this combined approach could clarify if these effects are attributed only to the strains we included here. Additionally, to overcome the resource limiting issues in viral research, we have established a basis for integrating computational immunology with experimental validation to advance understanding of ICV and IDV antigenicity.

## Acknowledgements

We thank Juggragarn Jengarn for his contribution to the experimental work.

## Author Contribution

**Conceptualisation:** MGM, SRK, CMT, NJT, KdC; **Formal Analysis:** MGM, SRK, KdC; **Data curation:** EM, NJT, CMT, KdC; **Investigation:** MGM, SRK, MMN, KdC; **Resources:** JD, PMD; HEE; EM, CMT, NJT, **Supervision:** EM, CMT, NJT, KdC; **Visualisation:** MGM, SRK; KdC; **Funding:** EM, CMT, NJT; **Writing – original draft preparation**: MGM; **Writing – review and editing:** all the authors.

## Bibliography

1. Krammer F, Smith GJD, Fouchier RAM, Peiris M, Kedzierska K, Doherty PC, Palese P, Shaw ML, Treanor J, Webster RG, García-Sastre A. Influenza. Nat Rev Dis Primers. 2018 Jun 28;4(1):3. doi: 10.1038/s41572-018-0002-y. PMID: 29955068; PMCID: PMC7097467.

2. Su, S.; Fu, X.; Li, G.; Kerlin, F.; Veit, M. Novel Influenza D virus: Epidemiology, pathology, evolution and biological characteristics. Virulence 2017, 8(8), 1580–1591. doi: 10.1080/21505594.2017.1365216.

3. Taylor, R. Studies on survival of Influenza virus between epidemics and antigenic variants of the virus. Am J Public Health Nations Health 1949, 39, 171–8.

4. Thielen, B.K.; Friedlander, H.; Bistodeau, S.; Shu, B.; Lynch, B.; Martin, K.; Bye, E.; Como-Sabetti, K.; Boxrud, D.; Strain, A.K.; Chaves, S.S.; Steffens, A.; Fowlkes, A.L.; Lindstrom, S.; Lynfield, R. Detection of Influenza C Viruses Among Outpatients and Patients Hospitalized for Severe Acute Respiratory Infection, Minnesota, 2013–2016. Clin Infect Dis 2018, 66(7), 1092–1098. 10.1093/cid/cix931.

5. Guo, Y.J.; Jin, F.G.; Wang, P.; Wang, M.; Zhu, J.M. Isolation of influenza C virus from pigs and experimental infection of pigs with influenza C virus. J Gen Virol 1983, 64(Pt 1), 177–82. PMID: 6296296.

6. Zhang, H.; Porter, E.; Lohman, M.; Lu, N.; Peddireddi, L.; Hanzlicek, G.; Marthaler, D.; Liu, X.; Bai, J. Influenza C virus in cattle with respiratory disease, United States, 2016–2018. Emerg Infect Dis 2018, 24, 1926–1929.

7. Manuguerra, J.C.; Hannoun, C. Natural infection of dogs by influenza C virus. Res Virol 1992, 143(3), 199–204. doi: 10.1016/s0923-2516(06)80104-4. PMID: 1325663.

8. Horimoto, T.; Gen, F.; Murakami, S.; Iwatsuki-Horimoto, K.; Kato, K.; Akashi, H.; Hisasue, M.; Sakaguchi, M.; Kawaoka, Y.; Maeda, K. Serological evidence of infection of dogs with human influenza viruses in Japan. Vet Rec 2014, 174, 96. doi: 10.1136/vr.101929.

9. Hause, B.M.; Ducatez, M.; Collin, E.A.; Ran, Z.; Liu, R. et al. Isolation of a novel swine influenza virus from Oklahoma in 2011 which is distantly related to human influenza C viruses. PLoS Pathog 2013, 9, e1003176.

10. Ducatez, M.F.; Pelletier, C.; Meyer, G. Influenza D virus in cattle, France, 2011–2014. Emerg Infect Dis 2015, 21, 368–71. doi: 10.3201/eid2102.141449. PMID: 25628038.

11. Gorin S, Richard G, Hervé S, et al. Characterization of Influenza D Virus Reassortant Strain in Swine from Mixed Pig and Beef Farm, France. Emerg Infect Dis. 2024;30(8):1672–1676. doi:10.3201/eid3008.240089.

12. Benito AA, Monteagudo LV, Lázaro-Gaspar S, Garza-Moreno L, Antón-Baltanás N, Quílez J. First Detection and Genetic Characterization of Influenza D Virus in Cattle in Spain. Vet Sci. 2026;13(2):130. Published 2026 Jan 29. doi:10.3390/vetsci13020130.

13. Ohira K, Yokoe K, Li K, et al. Seroprevalence of influenza C and D virus infections among cattle in Japan. Vet Anim Sci. 2025;29:100468. Published 2025 Jun 6. doi:10.1016/j.vas.2025.100468.

14. Ruiz M, Puig A, Bassols M, Fraile L, Armengol R. Influenza D Virus: A Review and Update of Its Role in Bovine Respiratory Syndrome. Viruses. 2022 Dec 5;14(12):2717. doi: 10.3390/v14122717. PMID: 36560721; PMCID: PMC9785601.

15. C. M. Trombetta, A. Stufano, V. Biagini, et al., “Seroprevalence of Influenza D Virus in Cattle Workers: An Occupational Health Perspective,” Journal of Medical Virology 98 (2026): 1–7, 10.1002/jmv.70817.

16. White, S.K.; Ma, W.; McDaniel, C.J.; Gray, G.C.; Lednicky, J.A. Serologic evidence of exposure to influenza D virus among persons with occupational contact with cattle. J. Clin. Virol. 2016, 81, 31–33. PMID: 27294672.

17. Gao H, Sun W, Lu P, et al. Efficient airborne transmission of influenza D virus in ferret models and serological evidence of human exposure in Northeast China. Emerg Microbes Infect. 2025;14(1):2564308. doi:10.1080/22221751.2025.2564308.

18. Shimizu K, Kawakami C, Matsuzaki Y, et al. Monitoring Influenza C and D Viruses in Patients With Respiratory Diseases in Japan, January 2018 to March 2023. Influenza Other Respir Viruses. 2024;18(6):e13345. doi:10.1111/irv.13345.

19. Song H, Qi J, Khedri Z, Diaz S, Yu H, et al. (2016) An Open Receptor-Binding Cavity of Hemagglutinin-Esterase-Fusion Glycoprotein from Newly-Identified Influenza D Virus: Basis for Its Broad Cell Tropism. PLOS Pathogens 12(1): e1005411. 10.1371/journal.ppat.1005411.

20. Wiley DC, Skehel JJ. The structure and function of the hemagglutinin membrane glycoprotein of influenza virus. Annu Rev Biochem. 1987;56:365–94. doi: 10.1146/annurev.bi.56.070187.002053. PMID: 3304138.

21. Skehel JJ, Wiley DC. Receptor binding and membrane fusion in virus entry: the influenza hemagglutinin. Annu Rev Biochem. 2000;69:531–69. doi: 10.1146/annurev.biochem.69.1.531. PMID: 10966468.

22. Nissly RH, Zaman N, Ibrahim PAS, McDaniel K, Lim L, Kiser JN, Bird I, Chothe SK, Bhushan GL, Vandegrift K, Neibergs HL, Kuchipudi SV. Influenza C and D viral load in cattle correlates with bovine respiratory disease (BRD): Emerging role of orthomyxoviruses in the pathogenesis of BRD. Virology. 2020 Dec;551:10–15. doi: 10.1016/j.virol.2020.08.014. Epub 2020 Sep 26. PMID: 33010670; PMCID: PMC7519714.

23. Ramírez-Salinas GL, García-Machorro J, Rojas-Hernández S, Campos-Rodríguez R, de Oca AC, Gomez MM, Luciano R, Zimic M, Correa-Basurto J. Bioinformatics design and experimental validation of influenza A virus multi-epitopes that induce neutralizing antibodies. Arch Virol. 2020 Apr;165(4):891–911. doi: 10.1007/s00705-020-04537-2. Epub 2020 Feb 14. PMID: 32060794; PMCID: PMC7222995.

24. Qamar M. Sheikh, Derek Gatherer, Pedro A Reche, Darren R. Flower, Towards the knowledge-based design of universal influenza epitope ensemble vaccines, Bioinformatics, Volume 32, Issue 21, November 2016, Pages 3233–3239, 10.1093/bioinformatics/btw399.

25. Marotta MG, Neto MM, Daly J, et al. Development and optimisation of influenza C and influenza D pseudotyped viruses. J Virol Methods. 2026;339:115243. doi:10.1016/j.jviromet.2025.115243.

26. Burley S.K. et al. (2019). RCSB Protein Data Bank (RCSB PDB): biological macromolecular structures enabling research and education in fundamental biology, biomedicine, biotechnology, and energy. Nucleic Acids Research, 47(D1), D464–D474. https://www.rcsb.org/.

27. Johnson M., Zaretskaya I., Raytselis Y., Merezhuk Y., McGinnis S., & Madden T.L. (2008). NCBI BLAST: a better web interface. Nucleic Acids Research, 36(Web Server issue), W5–W9. 10.1093/nar/gkn201.

28. Clifford JN, Høie MH, Deleuran S, Peters B, Nielsen M, Marcatili P. BepiPred-3.0: Improved B-cell epitope prediction using protein language models. Protein Sci. 2022;31(12):e4497. doi:10.1002/pro.4497.

29. The PyMOL Molecular Graphics System, Version 3.0 Schrödinger, LLC. (https://www.pymol.org/support.html#cite-pymol).

30. Yan Y., Tao H., He J., & Huang S-Y. (2020). The HDOCK server for integrated protein– protein docking. Nature Protocols, 15(5), 1829–1852. 10.1038/s41596-020-0312-y.

31. Ferrara F, Temperton N. Pseudotype Neutralization Assays: From Laboratory Bench to Data Analysis. Methods Protoc. 2018 Jan 22;1(1):8. doi: 10.3390/mps1010008. PMID: 31164554; PMCID: PMC6526431.

32. Del Rosario JMM, da Costa KAS, Asbach B, Ferrara F, Ferrari M, Wells DA, Mann GS, Ameh VO, Sabeta CT, Banyard AC, Kinsley R, Scott SD, Wagner R, Heeney JL, Carnell GW, Temperton NJ. Exploiting Pan Influenza A and Pan Influenza B Pseudotype Libraries for Efficient Vaccine Antigen Selection. Vaccines (Basel). 2021 Jul 5;9(7):741. doi: 10.3390/vaccines9070741. PMID: 34358157; PMCID: PMC8310092.

33. Herrler G, Klenk HD. Structure and function of the HEF glycoprotein of influenza C virus. Adv Virus Res. 1991;40:213–34. doi: 10.1016/s0065-3527(08)60280-8. PMID: 1957719; PMCID: PMC7131673.

34. Katayama M, Murakami S, Ishida H, et al. Antigenic commonality and divergence of hemagglutinin-esterase-fusion protein among influenza D virus lineages revealed using epitope mapping. J Virol. 2024;98(3):e0190823. doi:10.1128/jvi.01908-23.

35. Zhu S, Harriman K, Liu C, et al. Human Cases of Highly Pathogenic Avian Influenza A(H5N1) — California, September–December 2024. MMWR Morb Mortal Wkly Rep 2025;74:127–133. DOI: 10.15585/mmwr.mm7408a1.

36. O’Callaghan RJ, Gohd RS, Labat DD. Human antibody to influenza C virus: its age-related distribution and distinction from receptor analogs. Infect Immun. 1980 Nov;30(2):500–5. doi: 10.1128/iai.30.2.500-505.1980. PMID: 7439993; PMCID: PMC551340.

37. Morse, S. S., Mazet, J. A., Woolhouse, M., Parrish, C. R., Carroll, D., Karesh, W. B., … & Daszak, P. (2012). Prediction and prevention of the next pandemic zoonosis. The Lancet, 380(9857),1956–1965. 10.1016/S0140-6736(12)61684-5PMC4663196.

