## Supplementary information for "Pseudotyped virus-based platform and structural analysis reveal potential cross-reactivity sites between influenza C and D viruses"

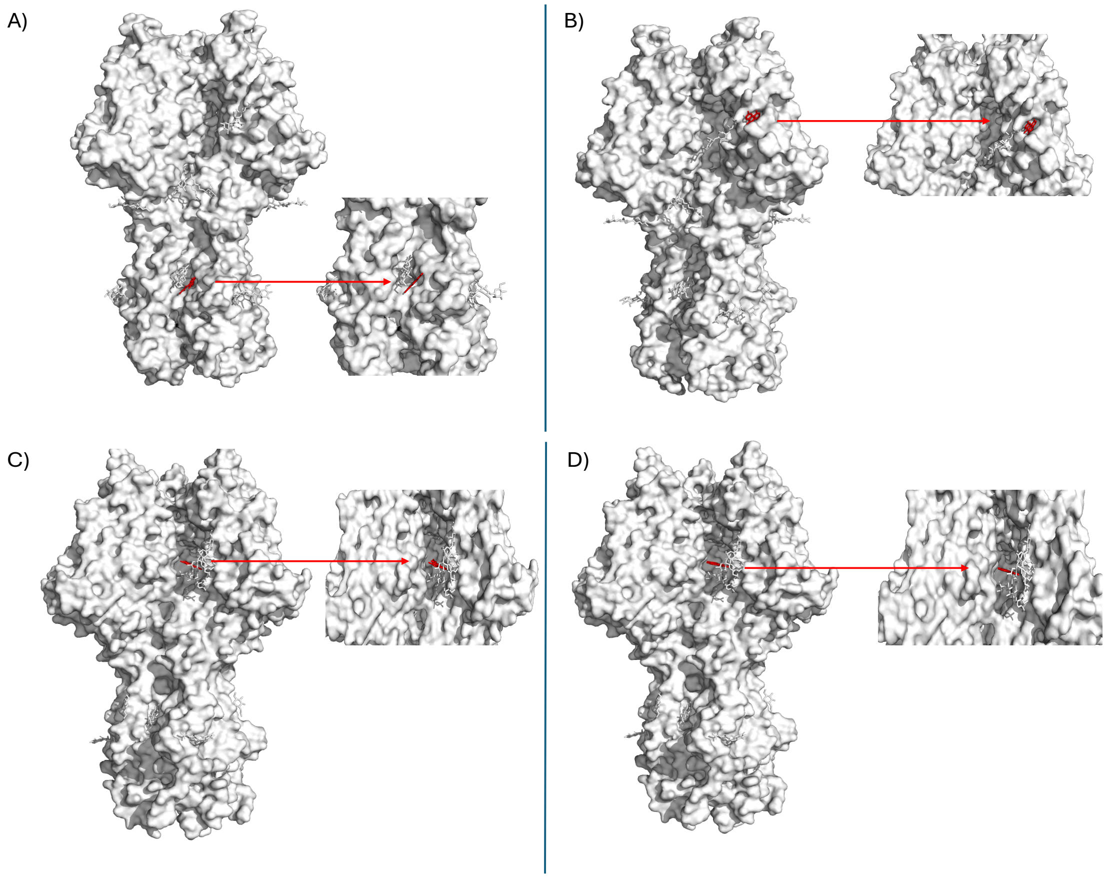


**Supplementary figure S1.** Molecular docking simulations of sialic acid receptors A) & C) Neu5Ac and B) & D) Neu5,9Ac with ICV HEF (A & B) and IDV HEF (C & D) using the HDOCK server. The sialic acids are shown in red, and the binding sites are indicated by red arrows.

```
C/Minnesota/33/2015
MFFSLLLVLGLTEAEKIKICLQKQVNSSFSLHNGFGGNLYATEEKRMFELVKPKAGASVL  60
D/swine/Italy/19974-3/2015
..LL.ATITAI.Q.KREL..IVQR..E.....S.....V.SMKTEP.TGFTNVTK....I  62

C/Minnesota/33/2015
NQSTWIGFGDSRTDKTNSAFPRSADVSAKTAGKFRSLSGGSLMLSMFGPPGKVDYLYQGC  120
D/swine/Italy/19974-3/2015
..KD..........LN.DQ..A.S..PLAV.K.......A.....A..............  122

C/Minnesota/33/2015
GKHKVFYEGVNWSPHAAIDCYSKNWTDIKLNFQKNIYELASKSHCMSLVNALDKTIPLHA  180
D/swine/Italy/19974-3/2015
..E..........E.G...FGS...QT.KD.YSK...A.RS.T..T...S..TK.STT.  183

C/Minnesota/33/2015
TAGTAKNCNNSFLKNPALYTQEVNPSEDKCGKENLAFFTLPTQFGTYECKLHLVASCYFI  240
D/swine/Italy/19974-3/2015
.....SS.SS.WM.S.LW.AES...GTKP..T.QS.T.....S..I.K.NK.V.QL...V  246

C/Minnesota/33/2015
YDSKEVYNKRGCDNYFQVIYDSSGKVVGGLDNRVSPYTGNSGDTPTMQCDMIQLKPGRYS  300
D/swine/Italy/19974-3/2015
.EN.TAF.TF..GD.Y.NY..GN.NLI..I....AA.R.IANVGVKIE.PSKI.N..T..  306

C/Minnesota/33/2015
VRSSPRFLLMPERSYCFDMKEKGPVTAIQSIWGKGRKSDYAVDQACLSTPGCMLIQKQRP  360
D/swine/Italy/19974-3/2015
I..T.....V.K......TDGGY.IQVV..E.SAS.R..N.TEE...Q.E..IF.K.TT.  366

C/Minnesota/33/2015
YIGEADDHHGDQEMRELLSGLDYEARCVSQSGWVNETSPFAEEYLLPPKFGRCPLAAKEE  420
D/swine/Italy/19974-3/2015
.V.....N...I...Q.....NNDTV......YTKGET..VRD..S...Y...Q.KTDSG  427

C/Minnesota/33/2015
SIPKIPDGLLIPTSGTDTTVTKPKSRIFGIDDLIIGLLFVAIVEAGIGGYLLGSRKESGG  480
D/swine/Italy/19974-3/2015
R..TL.S..I..QA...SLMRT.AT.........F...VGFVAGGVA...FW.RSNGG..  490

C/Minnesota/33/2015
GVTKESAEKGFEKIGNDIQILRSSTNIAIEKLNDRISHDEQAIRDLTLEIENARSEALLG  540
D/swine/Italy/19974-3/2015
.ASVS.TQA..D...K...Q..ND..A...GF.G..A......KN.AK...D..A...V.  550

C/Minnesota/33/2015
ELGIIRALLVGNISIGLQESLWELASEITNRAGDLAVEVSPGCWIIDDNICDQSCQNFIF  600
D/swine/Italy/19974-3/2015
......S.I.A.L.MN.K...Y...NQ..K.G.GI.Q.AG....YV.SEN..A..KEY..  610

C/Minnesota/33/2015
KFNETAPVPTIPPLDTKIDLQSDPFYWGSSLGLAITAAISLAALV  645
D/swine/Italy/19974-3/2015
N..GS.T...LR.V...VVIT...Y.L..TIA.CLLGLVAIV.S.  655
```

**Supplementary figure S2.** Amino acid sequence alignment between HEF glycoproteins of C/Minnesota/33/2015 (upper) and D/swine/Italy/19974-3/2015 (lower) strains, performed using NCBI Protein BLAST. Amino acids are referred to as residues according to structural biology conventions. Residue numbering follows the corresponding 3D models (PDB format), ensuring consistency between sequence annotations and structural representations. Dots represent conserved residues, while the capital letters indicate residue differences between the two sequences. Colored highlights indicate epitopes selected for mutant design. The red arrows indicate the locations of the site-specific substitutions introduced to generate the mutant constructs. Double-headed arrows indicate reciprocal exchanges of residues within shared epitopes, while single-headed arrows represent unidirectional substitutions in non-shared epitopes.

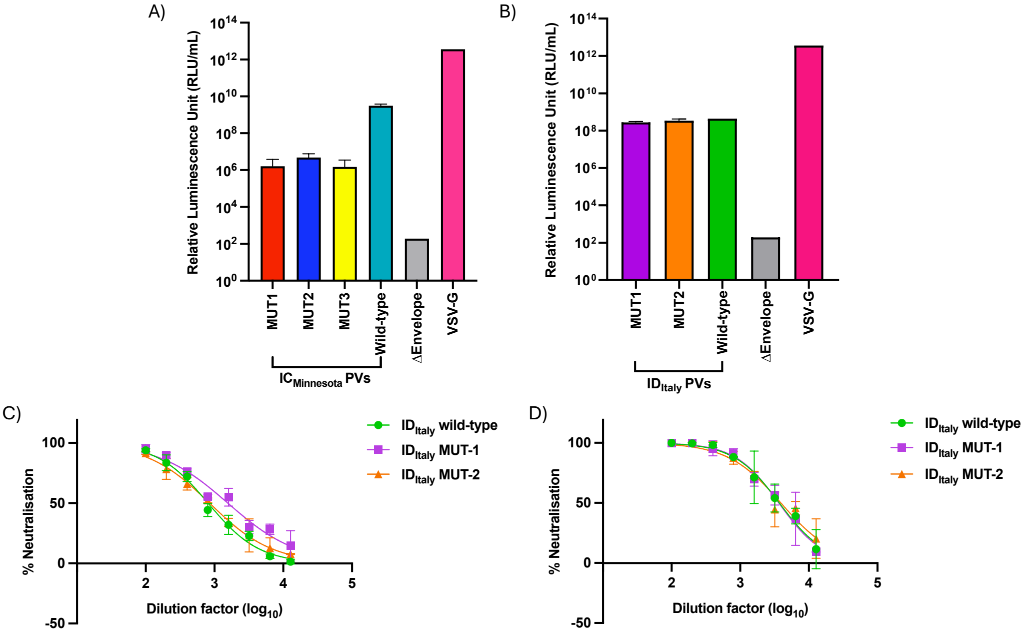


**Supplementary figure S3.** Titration of A) $IC_{Minnesota}$-mutant and wild-type PVs and B) $ID_{Italy}$-mutant and wild-type PVs. ST cells were transduced, and titres were measured as RLU/mL. The observed titre is compared with the positive control VSV-G PV and the negative control ΔEnvelope PV lacking a glycoprotein.
Neutralisation curves of $ID_{Italy}$ mutant and wild-type PVs of C) ICV antisera and D) IDV antisera. The y-axis represents the percentage of neutralisation across serum dilutions ($log_{10}$ scale) (x-axis). Data are shown as mean ± SD of duplicate wells.

**Supplementary table S1.** Summary of ICV and IDV HEF mutants generated. The table shows the targeted epitope, the corresponding color code used in the figures, the amino acid substitutions introduced in each mutant, and the number of residues exchanged between the two glycoproteins.

| Mutant name | Epitope | Amino acid substitutions | Number of residues exchanged |
|---|---|---|---|
| $HEF_{Minnesota}$-MUT1 | A (cyan) | K75L, T76N | 2 |
| $HEF_{Minnesota}$-MUT2 | D (yellow) | K334A, G335S, K337R | 3 |
| $HEF_{Minnesota}$-MUT3 | E (magenta) | H368N, Q372I | 2 |
| $HEF_{Italy}$-MUT1 | A (cyan) | L77K, N78T | 2 |
| $HEF_{Italy}$-MUT2 | E (yellow) | N374H, I378Q | 2 |
